# Oligomeric state of β-coronavirus non-structural protein 10 stimulators studied by OmniSEC and Small Angle X-ray Scattering

**DOI:** 10.1101/2023.06.26.546492

**Authors:** Wolfgang Knecht, S. Zoë Fisher, Jiaqi Lou, Céleste Sele, Shumeng Ma, Anna Andersson Rasmussen, Nikos Pinotsis, Frank Kozielski

## Abstract

Members of the β-coronavirus family such as SARS-CoV-2, SARS, and MERS have caused pandemics over the last 20 years. Future pandemics are likely and studying the coronavirus family members is necessary for their understanding and treatment. Coronaviruses possess 16 non-structural proteins, many of which are involved in viral replication and other vital functions. Non-structural protein 10 (nsp10) is an essential stimulator of nsp14 and nsp16, modulating RNA proofreading and viral RNA cap formation. Studying nsp10 of pathogenic coronaviruses is central to understanding its multifunctional role. We report the biochemical and biophysical characterisation of full-length nsp10 from MERS, SARS and SARS-CoV-2. Proteins were subjected to a combination of OmniSEC and SEC-MALS to characterise their oligomeric state. Full-length nsp10s were predominantly monomeric in solution, while truncated versions of nsp10 have a higher tendency to oligomerise. Small angle X-ray scattering (SAXS) experiments reveal a globular shape of nsp10 which is conserved in all three coronaviruses, including MERS nsp10, which diverges most from SARS and SARS-CoV-2 nsp10s. In conclusion, unbound nsp10 proteins from SARS, MERS, and SARS-CoV-2 are globular and predominantly monomeric in solution. Additionally, we describe for the first time a functional role of the C-terminus of nsp10 for tight binding to nsp14.

## Introduction

Coronaviruses (CoVs) are RNA viruses belonging to the Coronaviridae family, order Nidovirales. Within the family there are four genera named α, β, γ and σ coronaviruses. CoVs that generally infect humans are in the genera β-coronavirus, seven of which have been shown to affect the human host. Although four of those usually cause mild effects with symptoms like the common cold, the three others caused deadly outbreaks over the past 20 years. Of the three pathogenic CoVs, Severe Acute Respiratory Syndrome (SARS) caused a limited outbreak in China in 2002 with approximately 8000 infections and a mortality rate of close to 10% (World Health Organization, 2003; Petersen et al. 2020). 10 Years later a new CoV outbreak occurred in Saudi Arabia, named Middle East Respiratory Syndrome (MERS), which had a lower infection rate (ca. 2500) but a significantly higher mortality rate at ∼35% (World Health Organization, 2022).

The most recent CoV outbreak occurred towards the end of 2019 caused by a new CoV later named SARS-CoV-2 leading to Covid-19. SARS-CoV-2 caused a global, ongoing pandemic affecting the entire world population with over 670 million infections and more than 6.7 million deaths although the unofficial estimates are much higher (World Health Organization, 2023). It is now generally accepted that SARS-CoV-2 is soon becoming endemic in many countries (Torjesen, 2021). Due to the regular recurrence of CoVs over the last 20 years, it is expected that CoV-related outbreaks will occur in the future making it prudent to study and understand how CoVs infect humans, replicate in the human host, and how to handle infections to reduce severe disease and mortality.

Most scientific knowledge amassed about β-CoVs mostly originates from work carried out on SARS over the past 20 years. Many of those scientific insights are also relevant to SARS-CoV-2, which is not surprising given high sequence conservation in the coding regions of both genomes. SARS-CoV-2 is a single-stranded positive-sense RNA virus of approximately 30 kb coding for three distinct classes of proteins (Wu et al., 2022). Four structural proteins named spike, envelope, nucleocapsid and membrane form the virion. Seven so-called accessory proteins contribute to viral pathogenicity (Silvas et al., 2021) and may vary significantly between the various β-CoVs (Fang et al., 2021). The largest class of proteins is formed by the 16 non-structural proteins (nsp1 to nsp16), many of which are involved in the replication of the virus (Yadav et al., 2021) and are part of the replication-transcription complex (RTC).

Some of the nsps are validated drug targets with clinically used drugs such as Paxlovid (Nirmatrelvir and Ritonavir) targeting the main protease (nsp5) (Owen et al., 2021; Drug Administration, 2021) and Remdesivir inhibiting the RNA-dependent RNA polymerase (RdRp) complex (nsp12-nsp7 and nsp8) (Cotrim & Barros, 2022; Kokic et al., 2021). As their application is limited and the development of resistance is widely anticipated, the development of new drugs targeting other potential protein targets is highly warranted (Moghadasi et al., 2023; Gandhi et al., 2022). Recently, it has been shown in a clinical trial that approved monoclonal antibodies lost their effects against omicron variants and their use is therefore not recommended anymore (U.S. Food & Drug Administration, 2022; Wilhelm et al., 2022) clearly supporting the notion that the pandemic is not over yet.

Several other non-structural proteins are considered potential targets due to their essential roles in viral replication such as for example the helicase (nsp13), bifunctional nsp14 containing 3’-to-5’ exoribonuclease (ExoN) and N7 methyltransferase (N7-MTase) activities (Hsu et al., 2021), nd nsp16 with 2’-O-methyltransferase activity (2’-O-MTase) (Yadav et al., 2021; Decroly et al., 2011), although many other non-structural proteins are also investigated for their usefulness as potential targets.

One central nsp is nsp10, a scaffolding protein that interacts with and stimulates at least two other non-structural proteins, nsp14 and nsp16 (Viswanathan 2020; Krafcikova 2020). Nsp10 stimulates the ExoN activity of SARS and SARS-CoV-2 nsp14, but not its N7-MTase activity (Ma et al., 2015) by forming a 1:1 complex (Lin et al., 2021). Likewise, it also stimulates the 2’-O-MTase activity of nsp16, by forming a heterodimeric complex reported for all three pathogenic β-CoVs (Decroly 2011; Gupta 2021; Minasov 2020; PDB example entries 2XYR and 5YN8). This makes nsp10 a vital player not only in the RNA cap formation process (Chen 2016) but also for RNA proofreading (Shannon 2023). These proteins have been shown to methylate the N7 position of the cap guanyl (cap 0 formation), methylate the 2’ position of the ribose in the first base, and formation of the triphosphate bridge (cap 1 formation) resulting in viral RNA that is indistinguishable from host mRNA. Recently, it has even been proposed that nsp14, nsp10, and nsp16 can form a heterotrimeric complex, whereby nsp14 and nsp16 overlap on nsp10 in a particular N-terminal region of nsp14, termed the “lid” that shows high structural flexibility (Matsuda 2022, Czarna 2022). Successful targeting of nsp10 could lead to the inhibition of ExoN and N7-MTase functions, thus preventing RNA cap formation but also hindering RNA proofreading by the ExoN domain of falsely incorporated bases by the non-proofreading RdRp complex (Yan et al., 2021). Several knowledge gaps exist, such as the oligomeric state of the unbound form of nsp10 in solution compared to its state in forming its various heterodimeric complexes.

In this project, we characterise and compare the oligomeric state of nsp10 proteins from the SARS, MERS, and SARS-CoV-2 (Figure 1). The sequence alignment in Figure 1 illustrates the high degree of nsp10 sequence conservation of ∼62% for MERS compared to both SARS and SARS CoV-2, while nsp10 is 98% conserved between SARS and SARS-CoV-2. Figure 2 shows a structural overlay of available crystal structures of nsp10 from SARS, MERS, and SARS-CoV-2, indicating also the high degree of structural homology (r.m.s.d. is <1 Å^2^ over all atoms). Using various techniques, we show that these nsp10s are consistently and predominantly monomeric in solution. To further probe the shape, we also studied Small Angle X-ray Scattering (SAXS) data to confirm a conserved shape for nsp10s from SARS, MERS, and SARS-CoV-2 as being consistent with the known nsp10 crystal structures. Finally, we first time describe a functional role of the C-terminus of nsp10 for tight binding to nsp14 and that both proper N- and C-termini of nsp10 are important for optimal binding.

**Figure 1.**
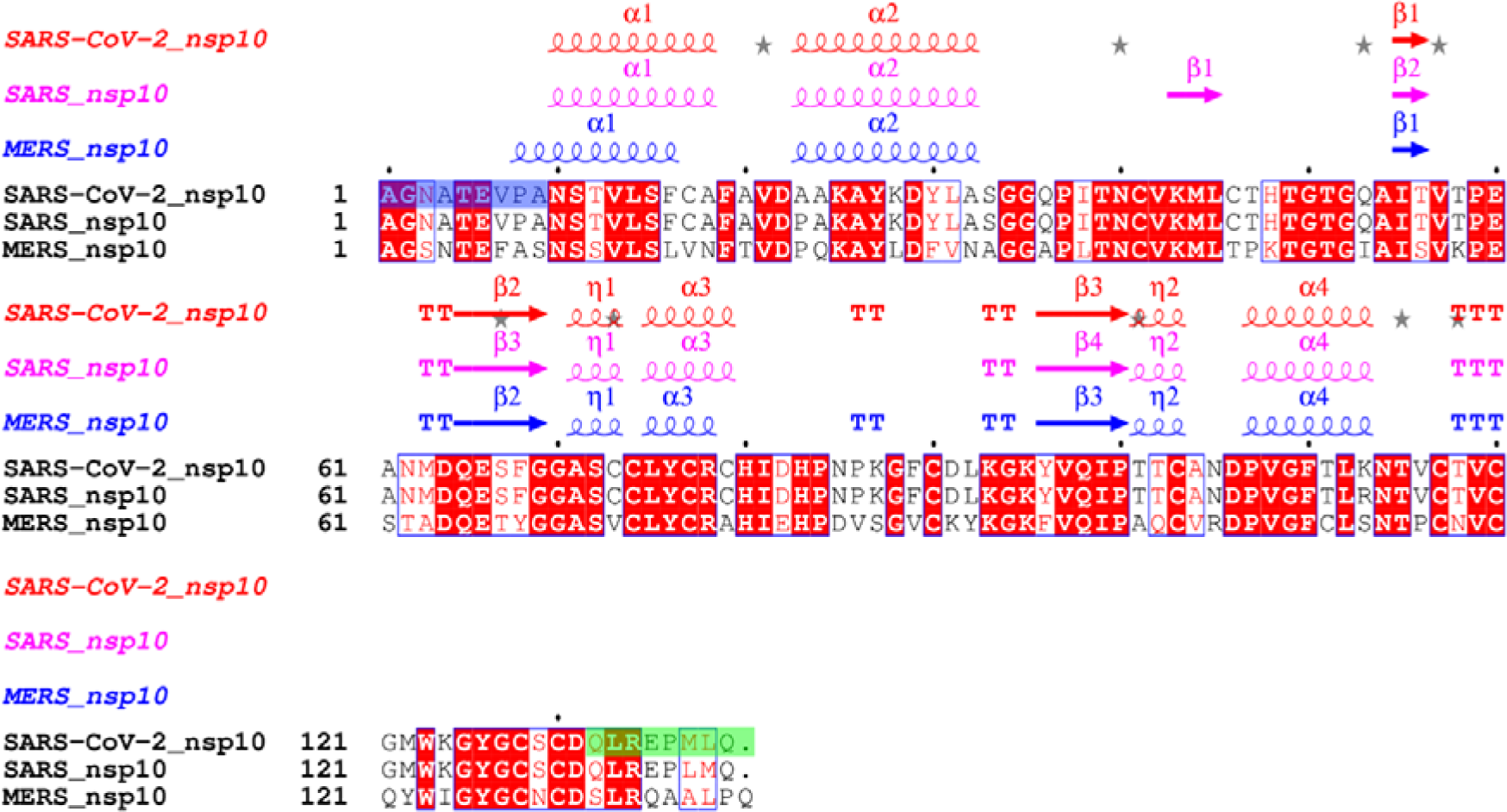
Protein sequence and structural alignment of full-length nsp10 proteins from SARS-CoV-2, SARS and MERS. Identical residues are shown in white on a red background, similar residues are shaded in red and distinct residues are displayed in black. Truncated residues of the long SARS-CoV-2 nsp10 construct are highlighted in green, and additional missing residues of the short SARS-CoV-2 nsp10 construct are highlighted in blue. The percent identity between SARS-CoV-2 and SARS nsp10 proteins is 97.1%. In contrast, the percent identity between SARS-CoV-2 and MERS nsp10 proteins is 59.4%. The alignment was carried out using CLUSTAL OMEGA, 1.2.4 (McWilliam 2013) while the secondary structure was assigned by dssp as implemented in ESPript 3.0 (Robert 2014).

**Figure 2.**
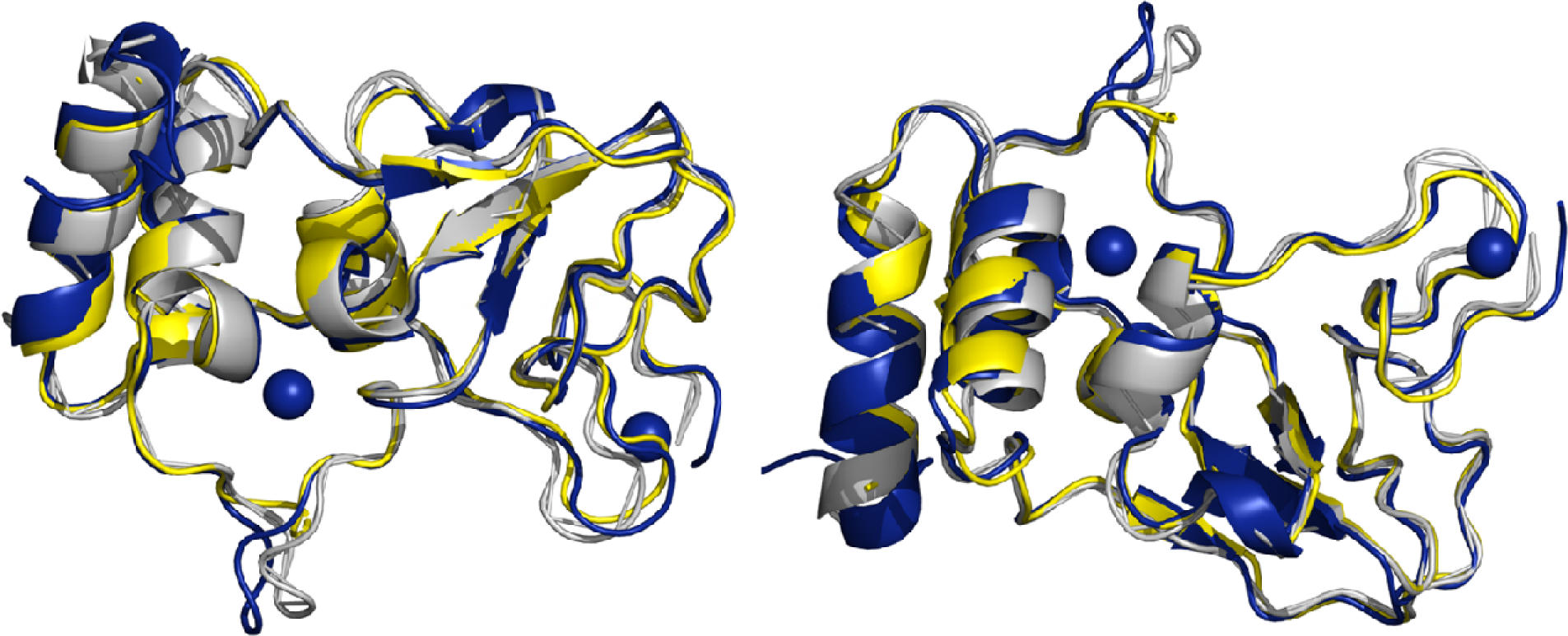
Overlay of crystal structures of nsp10 from MERS, SARS, and SARS-CoV-2. Ribbon diagram of crystal structures of nsp10 MERS (grey, PDB ID 5YN5), SARS (yellow, PDB ID 2G9T), and SARS-CoV-2 (blue, PDB ID 7DIY) with structural zinc atoms shown as blue spheres. View on right is rotated by 180∘ along horizontal axis with respect to view on the left.

## Materials and Methods

### Subcloning, expression and purification of beta-CoV nsp10 proteins

Expression constructs for nsp10 from SARS-CoV-2, SARS and MERS were synthesised at Genscript and subcloned into *E. coli* expression vector ppSUMO-2. Due to the cloning strategy, they contain up to two nonspecific residues (TM) at the N-terminus of the protein. Expressed proteins contain an N-terminal, cleavable His-SUMO double tag aiding in purification of the various proteins. The features of the expression clones are summarised in Table S1. A short SARS-CoV-2 nsp10 construct was designed as previously described (Rogstam 2020), covering residues Asn4264 (renumbered to Asn10 for convenience) to Gln4385, with a total length of 123 residues. This construct lacks residues at both the N- and C-termini. A long SARS-CoV-2 nsp10 construct consists of 133 residues and includes the N-terminus of the protein but is still devoid of the C-terminus. Full-length SARS-CoV-2 nsp10 contains 139 residues. Full-length SARS nsp10 is a protein of 139 residues. The protein sequence of SARS nsp10 (NP_828868, 139 residues), located between Ala4231 to Gln4369 of pp1a/pp1ab was used to generate the expression clone. MERS full-length nsp10 contains 140 residues.

The long and the short SARS-CoV-2 nsp10 proteins were expressed and purified as recently described (Rogstam 2020). Full-length SARS-CoV-2 nsp10 was expressed and purified as recently reported (Kozielski 2021). The full-length SARS and MERS nsp10s were purified using the same strategy employed for full-length SARS-Cov-2 nsp10. In brief, plasmids were transformed into Rosetta (DE3) competent cells (Merck KGaA, Darmstadt, Germany) and inoculated onto LB agar plates supplemented with 50 µg/ml kanamycin and 34 µg/ml chloramphenicol. For full-length MERS nsp10, the plasmid was transformed into BL21-Gold (DE3) competent cells (Agilent Technologies, Santa Clara, CA, USA) and inoculated onto LB agar plate supplemented with 50 ug/ml kanamycin.

A single colony from each LB agar culture was inoculated into 30 ml TB medium supplemented with the same antibiotics as above and incubated at 37°C and 220 RPM overnight. 5 ml of each culture was inoculated into each 1 L TB medium with the same antibiotics and incubated at 37°C and 220 RPM until the OD_600_ reached 0.6 - 1.0. Cultures were cooled down to 18°C and 1 mM IPTG was added to induce protein expression. The induced cultures were kept at 18°C, and 220 RPM for 20 - 24 h. After expression, cultures were centrifuged at 8,000 g at 4°C for 20 min to remove the medium. Cell pellets were resuspended in nsp10 buffer A (50 mM sodium phosphate buffer (NaPO_4_) pH 8.0, 300 mM NaCl, 20 mM Imidazole and 1 mM PMSF), flash-frozen in liquid nitrogen and stored at −80°C.

Cell pellets were thawed followed by sonication on ice for 10 cycles (30 sec on, 60 sec off) at a 16 mm amplitude, followed by centrifugation at 50,000 x g at 4°C for 1 h to remove cell debris. The supernatant was loaded into a 5 ml HisTrap FF crude column (Cytiva Life Sciences, Buckinghamshire, UK) pre-equilibrated with nsp10 buffer B (50 mM NaPO_4_ pH 8.0, 300 mM NaCl and 20 mM Imidazole). The loaded column was washed with 50 column volumes (CVs) nsp10 buffer B, followed by elution with 20 CVs nsp10 buffer C (50 mM NaPO_4_ pH 8.0, 300 mM NaCl and 250 mM Imidazole). Collected elution fractions were evaluated by SDS-PAGE and those containing nsp10 were pooled. ULP1 and 2 mM DTT were added to the pooled sample and dialysed against 2 L nsp10 dialysis buffer (50 mM NaPO_4_ pH 8.0, 300 mM NaCl, 20 mM Imidazole and 2 mM DTT) at 4°C overnight to cleave off the SUMO and His double tag from nsp10. Cleaved nsp10 was purified through 2 x 5 ml HisTrap FF crude columns pre-equilibrated with nsp10 buffer B to remove any uncleaved fusion protein, His-tagged ULP1 protease and other contaminations. Buffer exchanges were carried out by using 15 ml Amicon Ultra Centrifugal Units with 10,000 MWCO (Merck KGaA, Darmstadt, Germany).

### Purification of full-length nsp14

The expression and purification of full-length SARS-CoV-2 nsp14 was conducted as previously described (Kozielski 2021). It was expressed in *E. coli* TUNER (DE3) cells (Novagen, Darmstadt, Germany) using the following culture conditions: cells were grown in 9 L of LB Broth Miller (BD Difco) supplemented with 50 µg/ml kanamycin in 2.5 L Full-Baffle Tunair Shake Flasks (IBI Scientific, Dubuque, Iowa, USA), with 1 L/flask. Cells were grown at 18°C while shaking at 250 rpm. At OD_600_ of 0.9, the expression of nsp14 was induced by the addition of 0.01 mM IPTG, and growth continued for 20 h before the cells were harvested at 8000 × g for 20 min at 4°C.

Cell pellets were resuspended in 50 mM NaPO_4_, 300 mM NaCl and 20 mM imidazole at pH 8.0, supplemented with Complete Protease Inhibitor (EDTA free) (Roche, Stockholm, Sweden). The cells were lysed by passing the cell suspensions twice through a French pressure cell at 18,000 psi. Cell debris was removed by ultracentrifugation at 100,000× g for 60 min at 4°C and further clarified by filtration with a 0.45 µm filter cup. The clarified lysate was loaded onto a 5 mL HisTrap HP column (Cytiva, Uppsala, Sweden) connected to an ÄKTA Pure chromatography system (Cytiva, Uppsala, Sweden). The column was washed with 50 CVs of 50 mM NaPO4, 300 mM NaCl and 20 mM imidazole at pH 8.0, and the bound protein was eluted with a 0–100% gradient of 50 mM NaPO_4_, 300 mM NaCl and 500 mM imidazole at pH 8.0 over 20 CVs. Peak fractions were pooled and loaded onto a HiLoad 26/600 Superdex 75 pg column (Cytiva, Uppsala, Sweden) connected to an ÄKTA Purifier chromatography system (Cytiva, Uppsala, Sweden) with 50 mM HEPES, 150 mM NaCl and 1 mM DTT at pH 7.0 as the running buffer. Peak fractions containing pure nsp14 were pooled. The protein was concentrated using Amicon Ultra centrifugal filter units with 30 kDa MWCO (Millipore, Darmstadt, Germany). Protein concentrations were measured in a Nanodrop spectrophotometer using the theoretical extinction coefficients calculated using the Expasy ProtParam tool. The proteins were analysed on a Mini Protean Any kD TGX gel (BioRad, Solna, Sweden) stained with BioSafe Coomassie (BioRad, Solna, Sweden).

### MALDI-TOF/TOF mass spectrometry

SDS-PAGE gel bands were cut into 1×1 mm gel pieces and washed three times for 30 min by incubation with 400 µl 50% acetonitrile (ACN, Sigma-Aldrich) and 50 mM ammonium bicarbonate (ABC, Sigma-Aldrich). The gel pieces were dehydrated using 100% ACN before 25 µl of 12 ng/µl sequence grade modified trypsin porcine (Promega) in 25 mM ABC was added. The gel pieces in digestion buffer were incubated on ice for 3 h before further incubation overnight at 37°C. The following day Trifluoroacetic acid was added to a final concentration of 0.5% and peptides were extracted into a new tube ready for analysis by MALDI mass spectrometry.

MS and MS/MS spectra were acquired using an Autoflex Speed MALDI TOF/TOF mass spectrometer (Bruker Daltonics, Bremen, Germany) in positive reflector mode. Matrix solution, 0.5 µl consisting of 5 mg/ml α-cyano-4-hydroxy cinnamic acid, 80% acetonitrile, 0.1% TFA, was added to 1 µl peptide sample on a MALDI stainless steel plate. MS spectra were externally calibrated using Peptide Calibration Standard II (Bruker Daltonics) containing nine standard peptides (Bradykinin Fragment 1-7 (757.40), Angiotensin II (1046.54), Angiotensin I, m/z 1296.68; Substance P, m/z 1347.74; Bombesin m/z 1619.82; Renin Substrate,1758.93; ACTH clip 1-17, m/z 2093.09; ACTH clip 18-39, m/z 2465.20; Somatostatin 28, m/z 3147.47).

The identification of proteins was based on both MS and MS/MS data and was carried out with the Mascot Daemon software (version 2.4). The following search settings were used: trypsin as protease, two allowed missed cleavage sites, 50 ppm MS accuracy for peptides and 0.2 Da for MS/MS fragments, variable modifications: Oxidation (M). The files were searched against all organisms in SwissProt (updated 221008).

### OmniSEC experiments

The in-solution oligomeric state of the various nsp10 proteins was evaluated in running buffer composed of 50 mM Tris-HCl pH 8.0 and 150 mM NaCl. 100 µg of protein were injected in triplicate in an OMNISEC system (Malvern Panalytical, Malvern, UK) composed of the OMNISEC RESOLVE module (integrating a Superdex 75 Increase 10/300 GL column (Cytiva, Uppsala, Sweden), with a combined pump, degasser, autosampler and column oven) and the OMNISEC REVEAL, an integrated multi detector module (light scattering (RALS 90° angle and LALS 7° angle), differential refractive index, viscometer and diode-array-based UV/Vis spectrometer). The flow rate used was 0.5 ml/min and the detectors were normalised with bovine serum albumin (Thermo Fisher Scientific, Waltham, USA). The sample changer was cooled down to 4°C while the column oven and detector compartment were temperature controlled at 25°C. Data was collected and analyzed with the OMNISEC v11.32 integrated software provided by Malvern. The protein concentration was determined using an average refractive index increment (dn/dc) of 0.185 ml/g. Based on the reproducibility of the results, one injection from the triplicate was chosen for the calculations.

*SEC-MALS experiments.* The concentration for each protein was adjusted to 5 mg/ml. The samples were measured in an HPLC (Agilent 1100) in line connected to an eight-channel light scattering detector (Wyatt DAWN 8+) and a differential refractive index detector (Wyatt Optilab T-rex). The size-exclusion chromatography column (Superdex 75 10/300, Cytiva) was pre-equilibrated using buffer A (10 mM HEPES pH 7.5, 300 mM NaCl, 1.5% (v/v) Glycerol and 2 mM DTT) for 2 CVs and then 100 μL of each protein sample were injected using the auto-injector module of the HPLC at a flow rate of 0.5 ml/min. The main peaks were analysed using the ASTRA v6 software (Wyatt).

### Small Angle X-ray Scattering (SAXS)

After initial characterisation of nsp10s in solution by OmniSEC and SEC-MALS experiments, we used SAXS to study the overall shape and properties of various nsp10 proteins. The proteins were thawed on ice and injected into a size-exclusion chromatography column (Superdex 75 10/300, Cytiva) to remove aggregates and exchange buffers to 10 mM HEPES pH 7.5, 300 mM NaCl, 1.5% (v/v) Glycerol and 2 mM DTT. The proteins were then concentrated using centrifugal filters (Amicon Ultra 10k) and measured at synchrotron beamline B21 at Diamond Light Source, UK. Each sample was injected into the size-exclusion chromatography column (Superdex 75 10/300, Cytiva) linked to an HPLC system (Agilent 1200) and was subsequently measured in-line on the X-ray cell. Data were recorded on an Eiger detector under vacuum. The sample concentrations were adjusted to 10.0 mg/ml except for the short nsp10 protein construct measured at 8.0 mg/mL and full-length MERS nsp10 measured at 7.3 mg/mL and 50 μl of each sample were injected into the column. The scattering profiles for all samples indicated a clear separation between the major peak corresponding to the monomeric proteins and a small peak corresponding to an oligomeric form of the protein, similar to the SEC-MALS scattering profiles.

The data were processed using the program CHROMICS (Panjkovich 2017) where baseline (buffer) and peak (protein) were interactively selected based on the intensity peak and the calculated R_g_ values for each recorded point. All data were processed through the ATSAS suite (Manalastas-Cantos 2021) where the radius of gyration R_g_, D_max_, and excluded volume (Porod) were evaluated using standard procedures. The molecular masses of the solutes were evaluated from the Porod volume and also from the ratio of forward scattering I(0) of each sample; bovine service albumin was used as reference. The scattering from the high-resolution models was evaluated by CRYSOL (Svergun 1995) while the *ab-initio* modeling was calculated using DAMMIF (Franke 2009). Rigid body modeling was performed by BUNCH (Petoukhov 2005). Data collection statistics, parameters and additional details of the SAXS measurements are summarized in Table S2.

For comparing SAXS data of nsp10 constructs to published crystal structures, we chose the SARS-CoV-2 nsp10 structure bound to nsp14 ExoN domain (PDB entry 7DIY) by removing nsp14 and trimming the nsp10 to residues 6-128. This represents a conserved fold also found in the apo SARS-CoV2 nsp10 (PDB entry 6ZPE), but has a less ordered N-terminal region. In the structure of the SARS-CoV-2 nsp10 bound to nsp16, the N-terminal α-helix is disordered and at a completely different position compared to the two structures mentioned above.

### MST experiments to determine the binding affinity of nsp14 with various nsp10 constructs

Prior to the experiment, all nsp10 proteins were dialyzed (if needed) in 50 mM HEPES pH 7.0, 150 mM NaCl and 1 mM DTT buffer using 3.5 KDa MWCO Slide-A-Lyzer MINI dialysis buttons (ThermoFisher). His-SUMO-tagged full-length nsp14 was diluted to 200 nM with 50 mM HEPES pH 7.0, 150 mM NaCl, 0.05% Tween-20 and 1 mM DTT buffer and subsequently labelled by mixing with an equal volume of 100 nM RED-Tris-NTA labelling dye. The mix was then diluted twice, to a final concentration of 50 nM of labelled protein. The labelled protein solution was stored at 4 °C and kept away from light and always prepared fresh for each experiment. Serial 1:1 dilutions of nsp10 (200 µM starting concentration) were prepared in 50 mM HEPES pH 7.0, 150 mM NaCl, 0.05% Tween-20 and 1 mM DTT. An equal volume of 50 nM labelled full-length nsp14 was added to each diluted sample in the series, bringing the final concentration of labelled full-length nsp14 to 25 nM.

Samples were measured in a Monolith NT. 115 instrument (Nanotemper, München, Germany). The Nano-RED channel was used with 60 % excitation power and 60% MST power. The temperature control was kept at room temperature (22°C). Measurements were controlled by the Binding Affinity mode in MO.Control software and the data were analysed in ThermoAffinity (Burastero 2021).

## Results and Discussion

### Expression and purification nsp10 from β-CoV

We expressed nsp10 constructs of various lengths for MERS, SARS, and SARS-CoV-2. These include full-length proteins for all three CoVs, but also two shorter constructs for SARS-CoV-2. All proteins could be purified to an estimated purity of > 95% (Figure 3). On SDS-PAGE, full-length MERS nsp10 migrates to a position close to the lower band of each full-length SARS and SARS-CoV-2nsp10. The splitting of the protein band as observed on the SDS-PAGE (lane C and D) is not observed when the proteins are investigated in solution by OmniSEC (Table 1). The full-length MERS (lane B) and SARS-CoV-2 (lane D) nsp10 also showed a minor band at ca. 75 kDa, which was identified by mass spectrometry from the excised SDS-PAGE gel as the *E. coli* chaperone DNAK (Uniprot ID P0A6Y8, calculated MW of 69.1 kDa).

**Figure 3.**
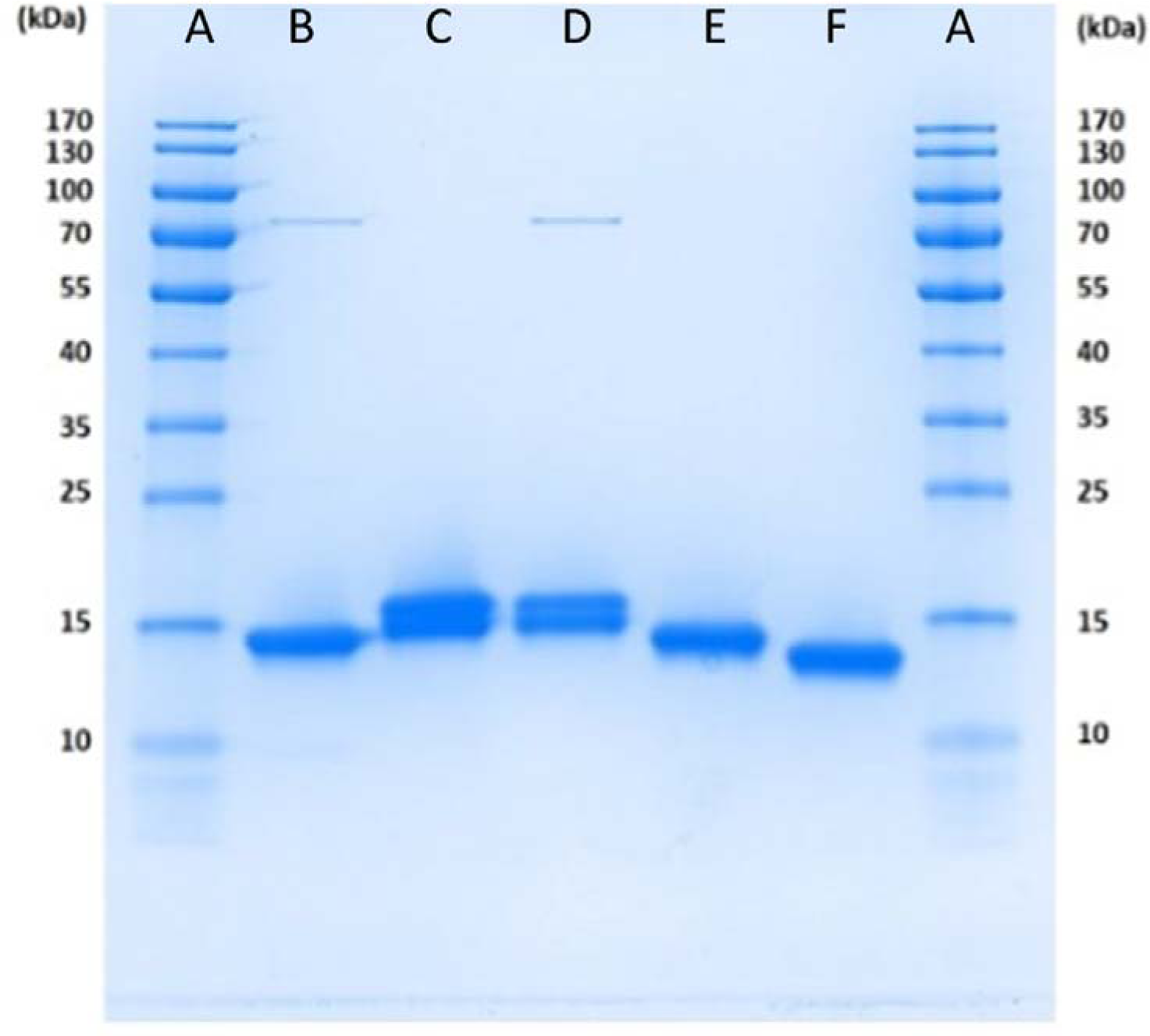
SDS-PAGE of SARS-CoV-2, SARS and MERS nsp10 constructs purified for this study. Lane designation: A) MW marker, B) full-length MERS nsp10, C) full-length SARS nsp10, D) full-length SARS-CoV-2 nsp10, E) long SARS-CoV-2 nsp10, F) short SARS-CoV-2 nsp10. The faint bands slightly above 70 kDa in (B) and (D) were identified by mass-spectrometry analysis as chaperone protein DNAK from *E. coli* (Uniprot ID: P0A6Y8; calculated MW: 69.1 kDa).

**Table 1.**
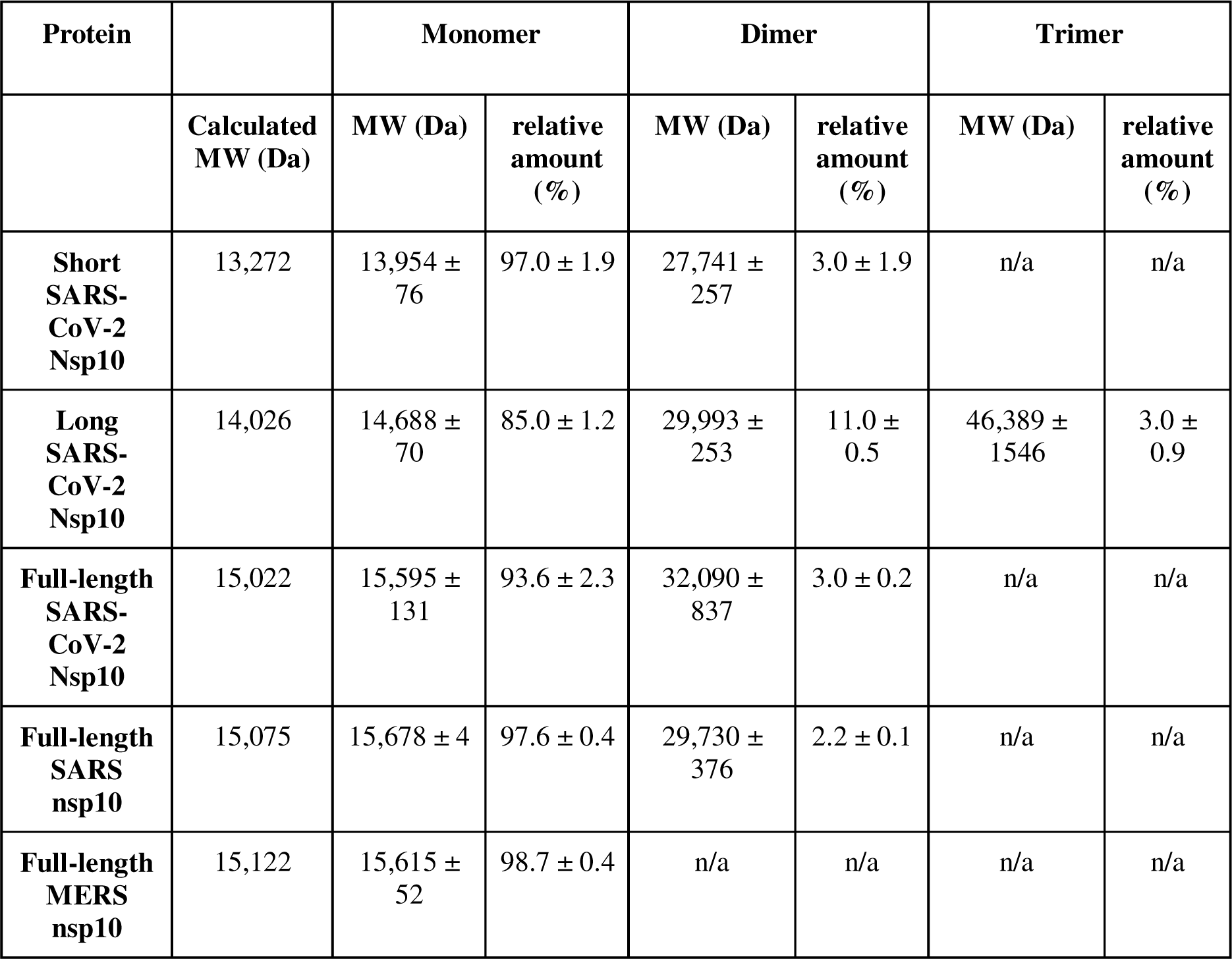
Molecular weights and oligomerisation of various nsp10s proteins determined by OmniSEC. OmniSEC runs with various nsp10 for the three β-CoVs. Only peaks with a relative amount of > 1.0 % are included in the table. MW and relative amount are average ± SD from three experiments.

| Protein |  | Monomer |  | Dimer |  | Trimer |  |
| --- | --- | --- | --- | --- | --- | --- | --- |
|  | Calculated MW (Da) | MW (Da) | relative amount (%) | MW (Da) | relative amount (%) | MW (Da) | relative amount (%) |
| **Short SARS-CoV-2 Nsp10** | 13,272 | 13,954 ± 76 | 97.0 ± 1.9 | 27,741 ± 257 | 3.0 ± 1.9 | n/a | n/a |
| **Long SARS-CoV-2 Nsp10** | 14,026 | 14,688 ± 70 | 85.0 ± 1.2 | 29,993 ± 253 | 11.0 ± 0.5 | 46,389 ± 1546 | 3.0 ± 0.9 |
| **Full-length SARS-CoV-2 Nsp10** | 15,022 | 15,595 ± 131 | 93.6 ± 2.3 | 32,090 ± 837 | 3.0 ± 0.2 | n/a | n/a |
| **Full-length SARS nsp10** | 15,075 | 15,678 ± 4 | 97.6 ± 0.4 | 29,730 ± 376 | 2.2 ± 0.1 | n/a | n/a |
| **Full-length MERS nsp10** | 15,122 | 15,615 ± 52 | 98.7 ± 0.4 | n/a | n/a | n/a | n/a |

### Determination of the oligomeric state of beta-CoV nsp10 proteins studied by OmniSEC and SEC-MALS

To determine the oligomeric state nsp10s from SARS, MERS, and SARS-CoV-2, the proteins were subjected to OmniSEC experiments. By combining the separation of the proteins by SEC and the information from light scattering at two different angles as well as the differential viscometer, the OmniSEC measurements allow reliable and highly sensitive measurements of the absolute molecular weight and distribution of the particles in solution. The OmniSEC results show that the molecular weight of all three full-length nsp10 in solution does not significantly differ from each other, indicating that the appearance of double bands in the SDS-PAGE (Figure 3) are likely to be artefacts. They also show that all constructs, in particular the full-length nsp10 proteins, are predominantly monomeric in solution (Table 1). Representative OmniSEC traces are shown in Figure S1. The full-length MERS nsp10 is particularly monodisperse, whereas full-length SARS and SARS-CoV-2 nsp10 form a minor pool of dimers (2 to 3%) in solution. A larger extent of dimer (3-11%) and trimer (up to 3%) formation can only be observed for the shorter, “artificial” SARS-CoV-2 protein constructs in solution. However, all three β-CoV nsp10s are predominantly monomeric in solution. SEC-MALS experiments conducted with the three SARS-CoV-2 protein constructs confirmed that nsp10 is predominantly monomeric with only a slight variation in the mass fraction (Figure S2) confirming our previous OmniSEC data.

Although the SARS nsp10 structure has previously been reported to be monomeric with one molecule in the asymmetric unit, it has been shown to be a dimer in solution using SEC (Joseph 2006). However, a truncated SARS nsp10 construct was used for size exclusion chromatography, not the full-length protein, as in our study. A fusion between SARS nsp10 and nsp11 has previously been reported to crystallise as a dodecamer (Su 2006). However, there is no evidence that such a nsp10-nsp11 fusion protein exists. In contrast, our novel results clearly demonstrate that full-length nsp10s from various medically relevant β-CoVs are predominantly monomeric and that truncated nsp10 constructs have a tendency to oligomerise into dimers and trimers, albeit to a very low extent. In conclusion, the nsp10s from β-CoV studied here are predominantly monomeric in solution in agreement with their functional roles as stimulators of nsp14 and nsp16 activities, acting in a 1:1 complex with their binding partners or as one partner in a nsp10-nsp14-nsp16 heterotrimeric complex.

### Small angle x-ray scattering reveals that nsp10s are compact monomers in solution

To gain further structural insights of the various nsp10 proteins in solution, we characterized the three SARS-CoV-2 nsp10 constructs (short, long, full-length) as well as the full-length SARS and MERS nsp10 proteins by small angle X-ray scattering using in-line SEC just before measuring SAXS data (SEC/SAXS). All scattering intensity plots display a similar profile to the one observed in SEC/MALS with a major peak and an almost negligent peak at an earlier elution volume, exemplified by the short and full-length SARS-CoV-2 nsp10 proteins (Figure S3).

The Radius of Gyration (R_g_) is consistent for all full-length proteins at about 17.2 - 17.3 Å and slightly smaller for the shorter SARS-CoV-2 nsp10 constructs. Consistently similar values were also calculated for the maximum distance (D_max_) ranging from 51 to 59 Å and for the estimated molecular masses. All statistics are summarised in Table S2. Overall SAXS reveals a similar shape of a globular domain for all full-length and shorter proteins (Figure 4). The distance distribution layout of the two shorter nsp10 proteins of SARS-CoV-2 resembles more closely a Gaussian layout, suggestive of a more spherical shape (Figure 4B). Kratky plots suggest a rigid shape for all measured samples (Figure S4).

**Figure 4.**
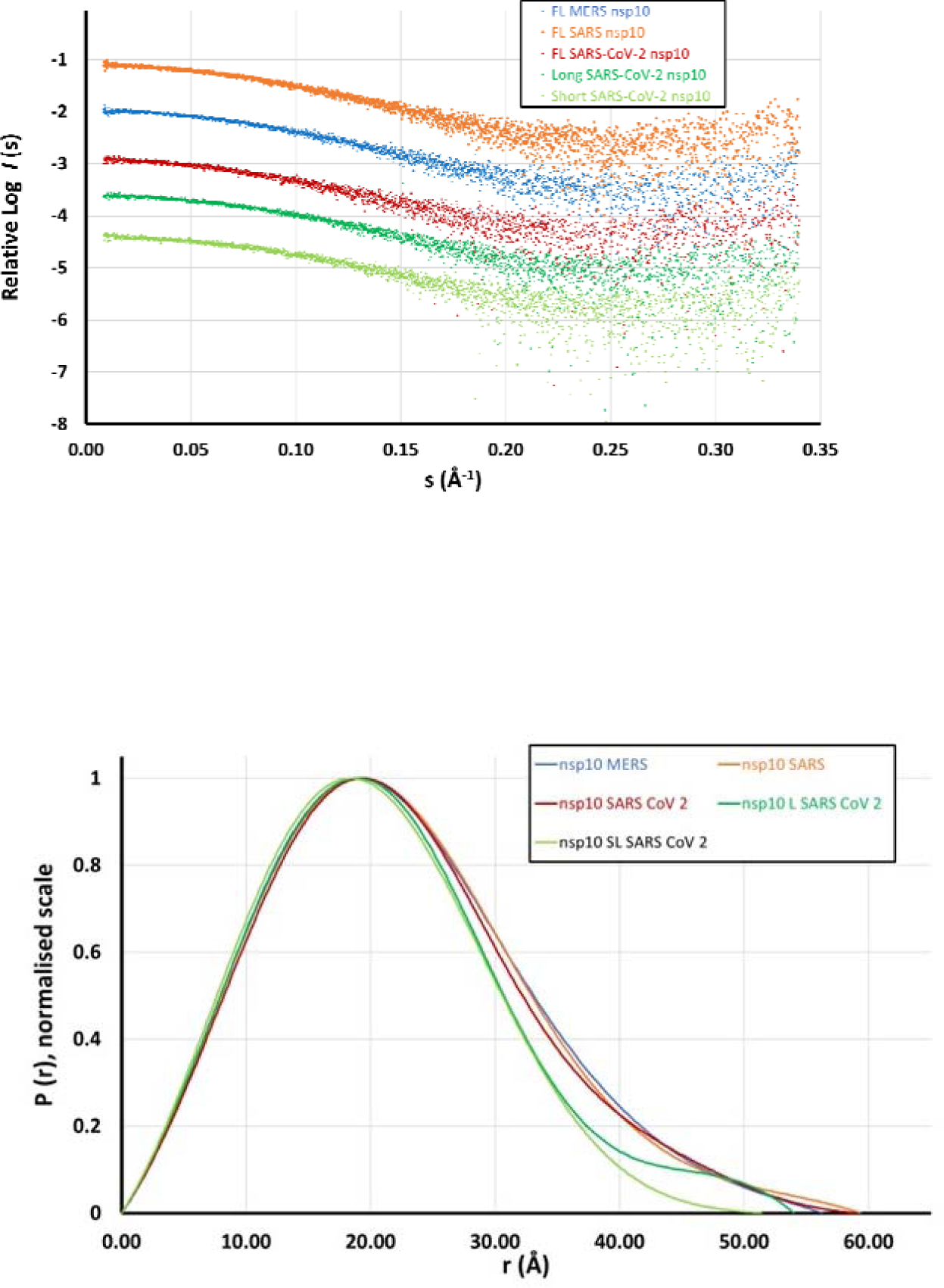
(A) Experimental SAXS data measured for nsp10 proteins from SARS-CoV-2 (short, long and full-length) and the full-length nsp10 proteins from MERS and SARS. Data are shown in relative scales. (B) Distance distribution profiles of the scattering data from panel A.

We further compared the scattering data for all five samples to a reference PDB. We selected the SARS-CoV-2 nsp10 coordinates from its crystal structure in complex with the nsp14-ExoN (PDB entry 7DIY) and trimmed the structure to nsp10 residues 6-128 as the most representative of the domain fold. The discrepancy χ^2^ as calculated from CRYSOL is ranging from 1.072 up to 1.252 for the four measured proteins and 2.408 for the long nsp10 SARS-CoV-2 construct. These values suggest a perfect match between the solution scattering data and the structure of nsp10 in the crystal (Figure 5).

**Figure 5.**
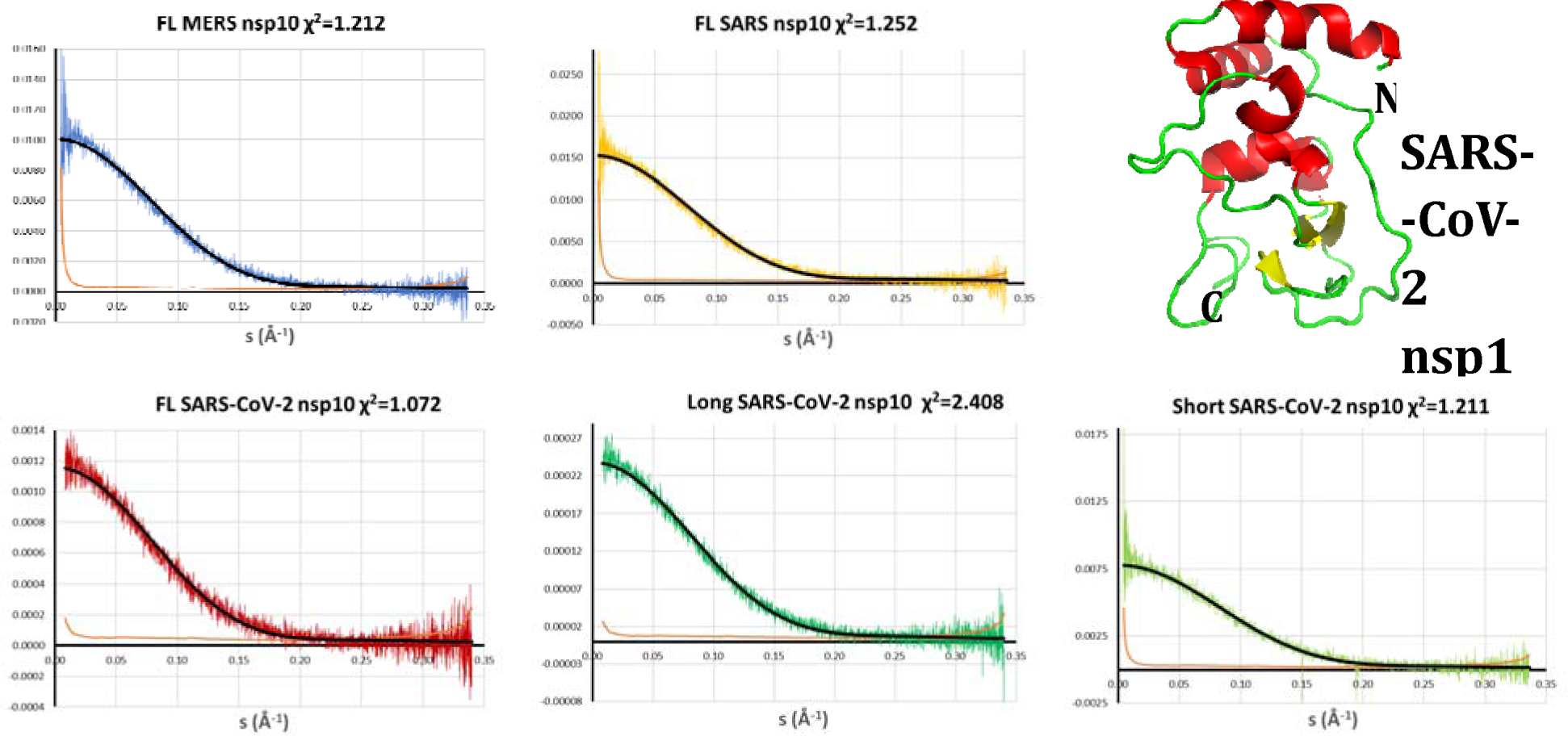
Comparison between the measured scattering data and the calculated scattering curve from the model (in black and in cartoon representation on the right). The green continuous line near the X-axis denotes the variation between experimental and calculated scattering data. Discrepancy χ^2^ is shown on the header of each plot. The nsp10 structure was extracted from the PDB entry 7DIY. FL full-length

To better understand the role of the residues at the N- and C-terminal regions of nsp10 we generated a low-resolution model for the full-length SARS-CoV-2 nsp10. We used a combined approach where we initially generated 20 *ab-initio* models using the program Dammif (Franke 2009). The models were aligned and an average envelope of them was used as the starting search volume for a slow mode *ab-initio* calculation using the software Dammin (Svergun 1999). The final model converged with a discrepancy χ^2^ against the raw data of 0.9876. We independently calculated a rigid body/*ab-initio* approach to fit the position of the missing residues at both termini using the software BUNCH (Petoukhov 2005). The final model reported a discrepancy χ^2^ of 1.01897 against the raw data, indicating a similarly excellent fit to the *ab initio* model. Superposition of the two models show a perfect fit where the termini appear to contribute to the compact core of the structure (Figure 6). This result is in agreement with the observations from the Kratky plots that nsp10 forms a rigid and stable protein domain. Very similar *ab-initio* / rigid body-refined models were obtained for full-length SARS and MERS nsp10 proteins.

**Figure 6.**
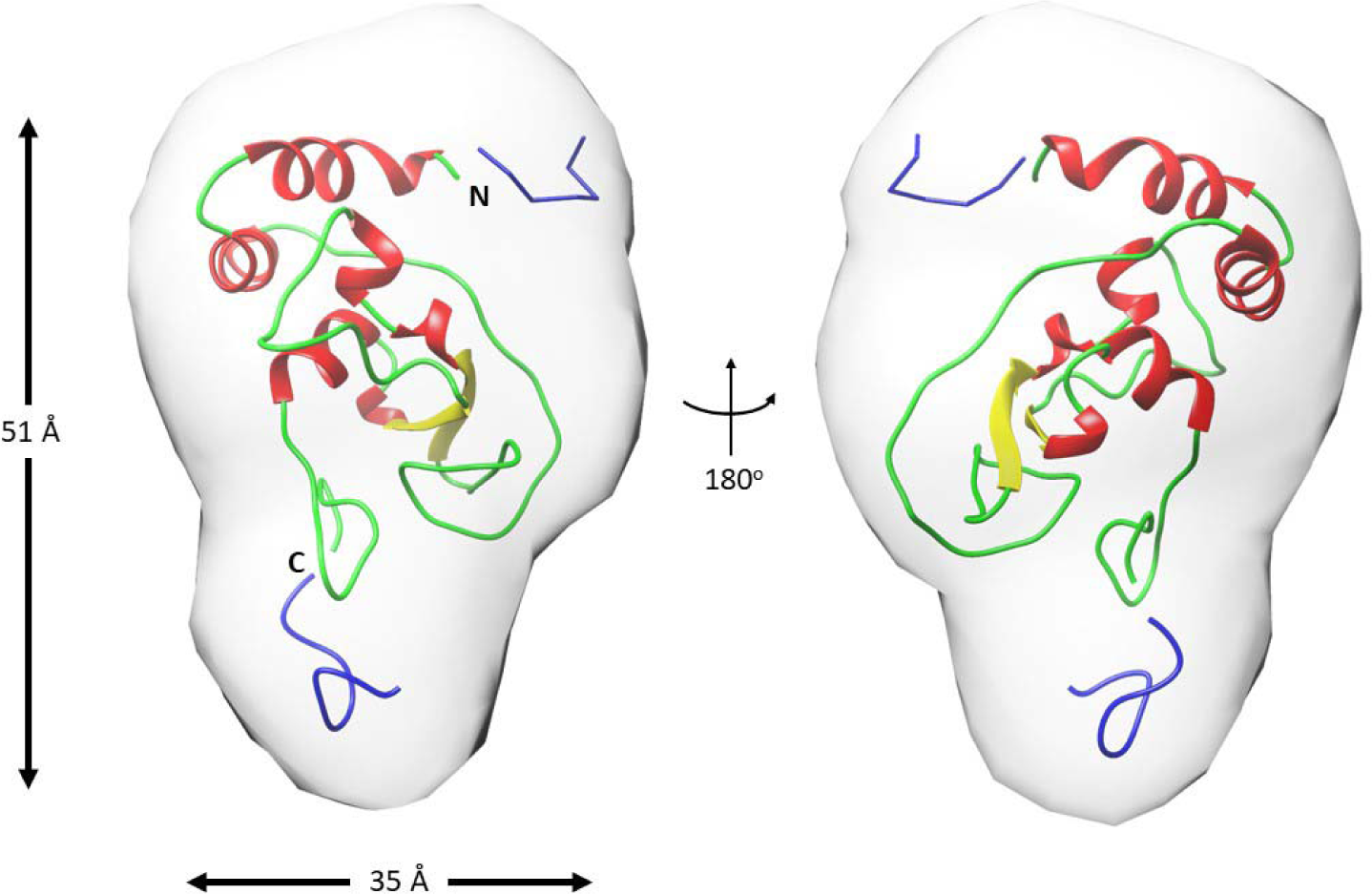
Superimposition of the full-length *ab-initio* nsp10 SARS-CoV-2 model shown as a molecular envelope with the rigid body / *ab-initio* model of the same protein shown as cartoon in two different orientations. The *ab-initio* envelope was generated in chimera using a map covering the calculated coordinates with a resolution of 13 Å. The extra residues added in the rigid body/*ab-initio* are shown in blue. Maximum dimensions for two of the orientations are indicated on the left as well as the N- and C-terminal boundaries of the model.

Overall, all nsp10 proteins from SARS-CoV-2, SARS, and MERS show an almost identical low-resolution shape. While this is expected for the SARS-CoV-2 and SARS nsp10 proteins where the sequence identity is 97%, it is less obvious for the MERS nsp10 which is only 60% identical to SARS-Cov-2 nsp10. Furthermore, the two full-length SARS-CoV-2 nsp10 models (*ab initio* and rigid body), combined with the evaluation of the data by CRYSOL using a SARS-CoV-2 nsp10 crystal structure, strongly suggest that extra residues at the termini of all proteins do not affect the overall fold and shape of the proteins.

### The affinity between nsp10 and nsp14 depends on the length of nsp10 and is influenced by an intact C-terminus

To better understand the role of the residues at the N- and C-terminal regions of nsp10 we also measured the affinity between nsp10 and nsp14. Whereas crystal structures of unbound nsp10, in the absence of their binding partners, possess a flexible and therefore not visible N-terminal region, this region becomes structured upon binding to either nsp14 or nsp16. To investigate the influence of the short N-terminal region of nsp10 on the affinity for its binding partner nsp14 we measured the binding affinity of the truncated and full-length versions of SARS-CoV-2 nsp10 using MST. The affinity for full-length nsp10 showed a K_d_ value of 0.74 ± 0.25 μM, whereas the short nsp10 construct lacking both N- and C-terminal regions showed a K_d_ value of 13.9 ± 1.6 μM, a 19-fold difference. Surprisingly, the long nsp10 construct lacking only a part of the C-terminus displays very weak affinity to nsp14 with a K_d_ value > than 100 μM. This result is unexpected because an interaction between the C-terminus of nsp10 and nsp14 is not even present in the crystal structures of either nsp10-nsp14 ExoN (SARS-CoV-2; PDB entry 7DIY)) or nsp10-nsp14 (SARS-CoV; 5NFY) complexes. These crystal structures do however not exclude an intermediate role of the C-terminus in achieving best binding between nsp10 and nsp14. These results show that the rigid core structure of nsp10 is sufficient for binding to nsp14 but that having both the proper N- and C-termini are important for tight or optimal binding. Furthermore, to the best of our knowledge this is the first time that a functional role for the C-terminus of nsp10 has been described.

## Abbreviations

CoV: Coronavirus
Covid-19: Coronavirus Disease 2019
DTT: DL-Dithiothreitol
ExoN: 3’-to-5’ exoribonuclease
IPTG: Isopropyl-β-D-thiogalactopyranoside
MALS: Multi Angle Light Scattering
MERS: Middle East Respiratory Syndrome
MST: MicroScale Thermophoresis
NaPO_4_: Sodium phosphate buffer
Nsp: non-structural protein
N7-MTase: N7 methyltransferase
PMSF: Phenylmethylsulfonylfluorid
RdRp: RNA-dependent RNA polymerase
SARS: Severe Acute Respiratory Syndrome
SARS-CoV-2: Severe Acute Respiratory Syndrome Coronavirus 2
SAXS: Small Angle X-ray Scattering
SEC: Size-Exclusion Chromatography

## Acknowledgements

The authors would like to thank Diamond Light Source for beamtime (proposal mx30393), and the staff of the beamline B21 for assistance with data collection. We are grateful to the MRC - UCL Therapeutic Acceleration Support (TAS) and the MRC DPFS (MR/X013995/1) for financial support. We are grateful to Drs. Tobias Schrader and Dominic Hayward (Ju[lich Centre for Neutron Science) for initial discussions on the project. We would also like to thank all Lund Protein Production Platform (LP3) staff for providing technical support. This research was initiated by the European Spallation Source ERIC for projects related to COVID-19 as well as Lund University and its faculties in their support of the Lund Protein Production Platform (LP3) and supported by grants from the Royal Physiographic Society of Lund and the Erik Philip-Sörensen Foundation. We acknowledge Katja Bernfur, Center for Molecular Protein Science, Lund University for performing the mass spectrometry analysis.

