## Supplementary data file for "Oligomeric state of β-coronavirus non-structural protein 10 stimulators studied by OmniSEC and Small Angle X-ray Scattering"

**Affiliation:** ^1^Department of Biology & Lund Protein Production Platform & Protein Production Sweden, Lund University, Sölvegatan 35, 22362 Lund, Sweden; ^2^European Spallation Source ERIC, P.O. Box 176, 22100 Lund, Sweden; ^3^School of Pharmacy, University College London, London, 29-39 Brunswick Square, London WC1N 1AX, United Kingdom; ^4^Institute of Structural and Molecular Biology, Birkbeck College, WC1E 7HX London, United Kingdom.

**Contents**

**Table S1**	Summary of nsp10 constructs from β-CoVs	2

**Table S2**	SAXS data collection and scattering parameters	3

**Fig S1**	Representative OmniSEC traces	4

**Fig S2**	SEC-MALS analysis of three SARS-CoV-2 nsp10 constructs.	5

**Fig S3**	Scattering profiles for two SARS-CoV- 2 nsp10 samples	6

**Fig S4**	Kratky plots for the nsp10 proteins measured by SAXS	7

**Table S1.** Summary of nsp10 constructs from β-CoVs used in this study.

| **Protein construct and name** | **Residue numbering** | **Original polypetide numbering** | **Calculated MW after cleavage [Da]** | **Calculated Ip after cleavage** |
| --- | --- | --- | --- | --- |
| **Short SARS-CoV-2 nsp10** | 10-133 | Asn4264 - Gln4385 | 13,272 | 7.70 |
| **Long SARS-CoV-2 nsp10** | 1-133 | Ala4255 - Gln4385 | 14,026 | 6.70 |
| **Full-length SARS-CoV-2 nsp10** | 1-139 | Ala4255 - Gln4391 | 15,022 | 6.70 |
| **Full-length MERS nsp10** | 1-140 | Ala4238 - Gln4378 | 15,122 | 7.70 |
| **Full-length SARS nsp10** | 1-139 | Ala4231- Gln4369 | 15,075 | 6.70 |

**Table S2.** SAXS data collection and scattering parameters for the β-CoV nsp10 proteins studied.

| **Sample** | **SARS-CoV-2** | | | **SARS** | **MERS** |
| --- | --- | --- | --- | --- | --- |
|  | **full-length nsp10** | **long nsp10** | **short nsp10** | **full-length nsp10** | **full-length nsp10** |
| **Data Collection parameters** | | | | | |
| Instrument | B21 Beamline (Diamond Light Source, UK) | | | | |
| Beam size at sample (mm^2^) | 1.0 ´ 0.25 | | | | |
| Wavelength (Å) | 0.9464 | | | | |
| s-Range (Å^-1^) | 0.0045-0.3400 | | | | |
| Method | SEC-SAXS | | | | |
| Sample to Detector D (mm) | 3722.0 | | | | |
| Temperature (K) | 288 | | | | |
| **Structural parameters** | | | | | |
| *R_g_* (Å) (from Guinier) | 17.10  (± 0.14) | 16.35  (± 0.10) | 15.20  (± 0.12) | 17.19  (± 0.10) | 17.27  (± 0.10) |
| *R_g_* (Å) (from P(r)) | 17.31  (± 0.01) | 16.49  (± 0.01) | 15.46  (± 0.01) | 17.40  (± 0.02) | 17.32  (± 0.01) |
| *D_max_* (Å) | 59.0 | 54.0 | 51.4 | 59.4 | 53.3 |
| Porod volume estimate (Å^3^)* | 26542 | 23162 | 20976 | 24726 | 24363 |
| **Molecular mass determination** | | | | | |
| Molecular mass from Porod volume (Vp ∗ 0.6) (Da) | 15925 | 13897 | 12585 | 14836 | 14618 |
| Molecular mass from forward scattering (Da) | 15395  (± 268) | 13985  (± 291) | 9324  (± 161) | 20841  (± 63) | 14328  (± 44) |
| Molecular mass from sequence (Da) | 15151 | 14026 | 13272 | 14843 | 14891 |
| **Software** | | | |  |  |
| Data processing | ATSAS (PRIMUS, GNOM) | | | | |
| *Ab initio* analysis | DAMMIF | | | | |
| Atomic models scattering | CRYSOL | | | | |

*Porod Volume was calculated through Dammif.

**Figure S1.** Representative OmniSEC traces. **A.** Short SARS-CoV-2 nsp10. **B.** Long SARS-CoV-2 nsp10. **C.** Full-length SARS-CoV-2 nsp10. **D.** Full-length SARS nsp10. **E.** Full-length MERS nsp10. The refractive index is shown in red, the determined molecular weight of the peak is in green.

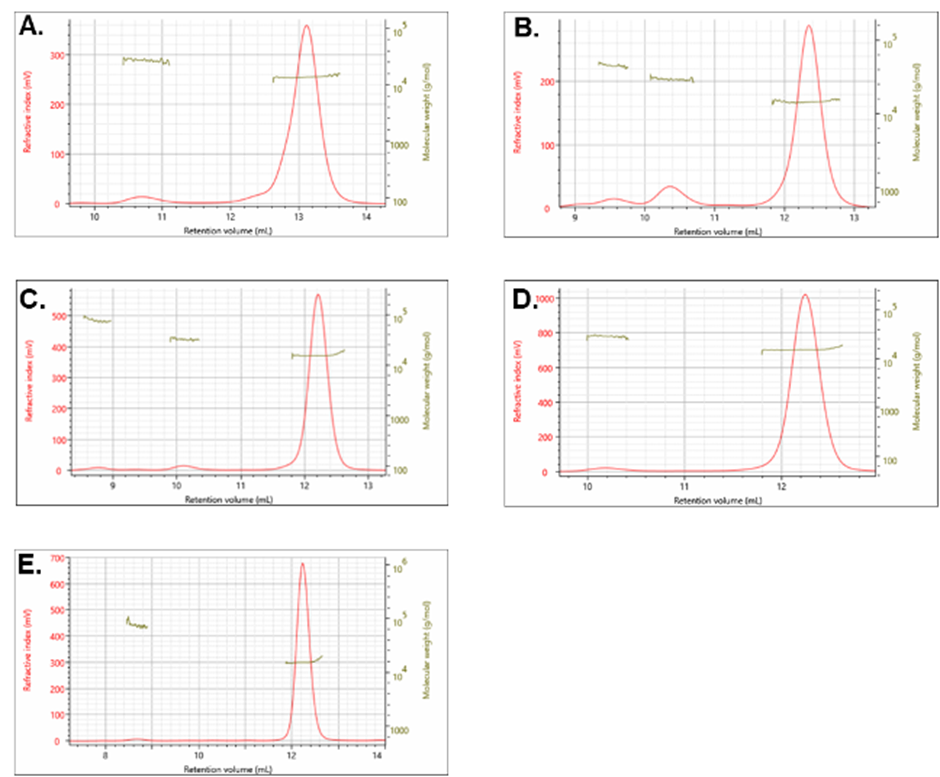

**Figure S2.** SEC-MALS analysis of three SARS-CoV-2 nsp10 constructs. Light scattering (LS) peaks of short, long, and full-length SARS-CoV-2 nsp10s are coloured in light green, dark gree, and red, respectively. The inset report on the molecular weight and mass fractions measured experimentally using SEC-MALS, where the lower peaks are categorised as peak 2 and the main peaks are named peak 2. The molecular weights for the second peak of short SARS-CoV-2 nsp10 was not statistically significant and therefore not included in the table.

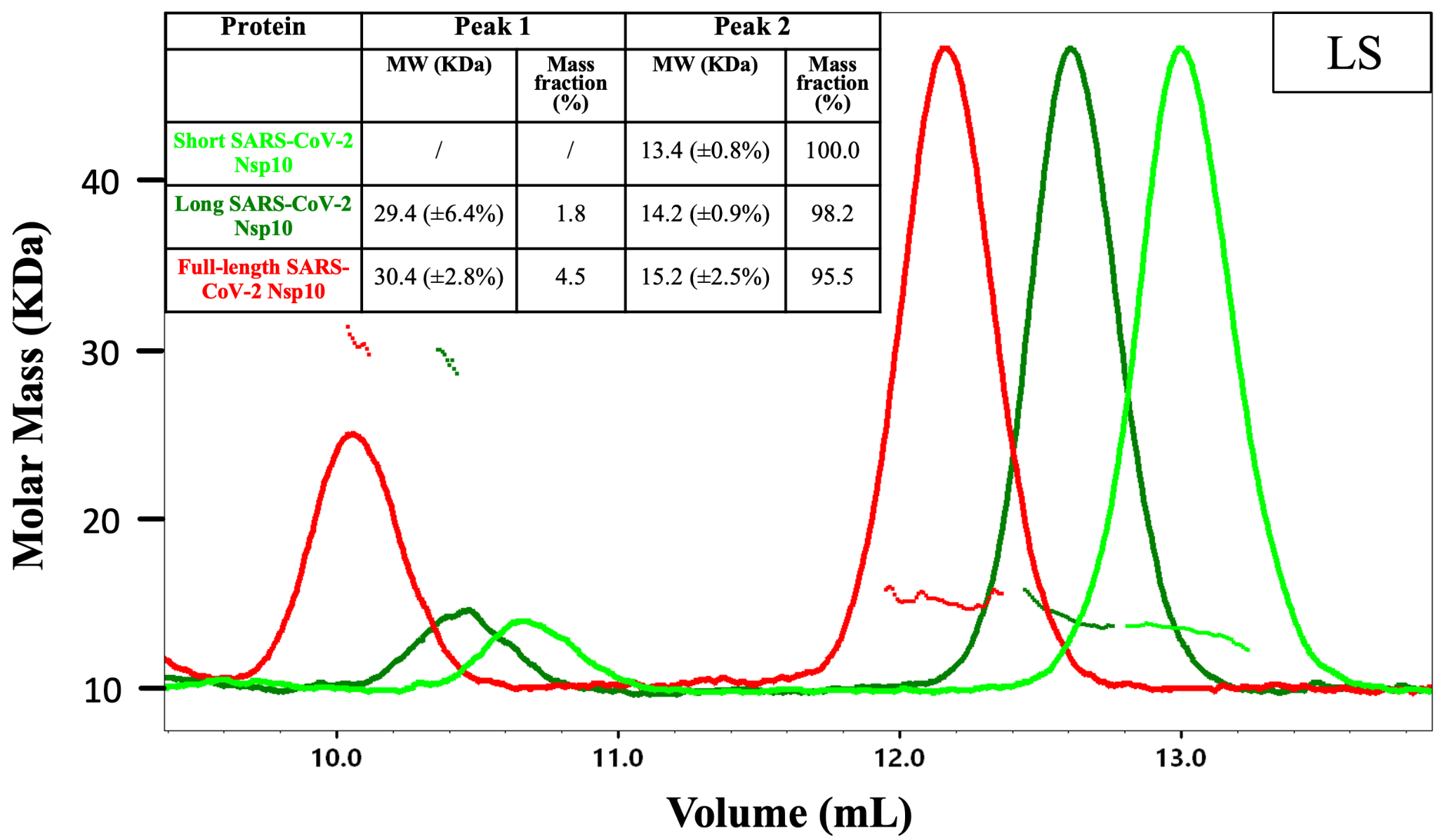

**Figure S3.** Scattering profiles for two short and full-length SARS-CoV-2 nsp10 samples, measured by SEC/SAXS. We recorded one frame for each 0.026 ml, which corresponds to an elution volume of 12.42 ml for full-length nsp10 and 13.45 ml for short nsp10.

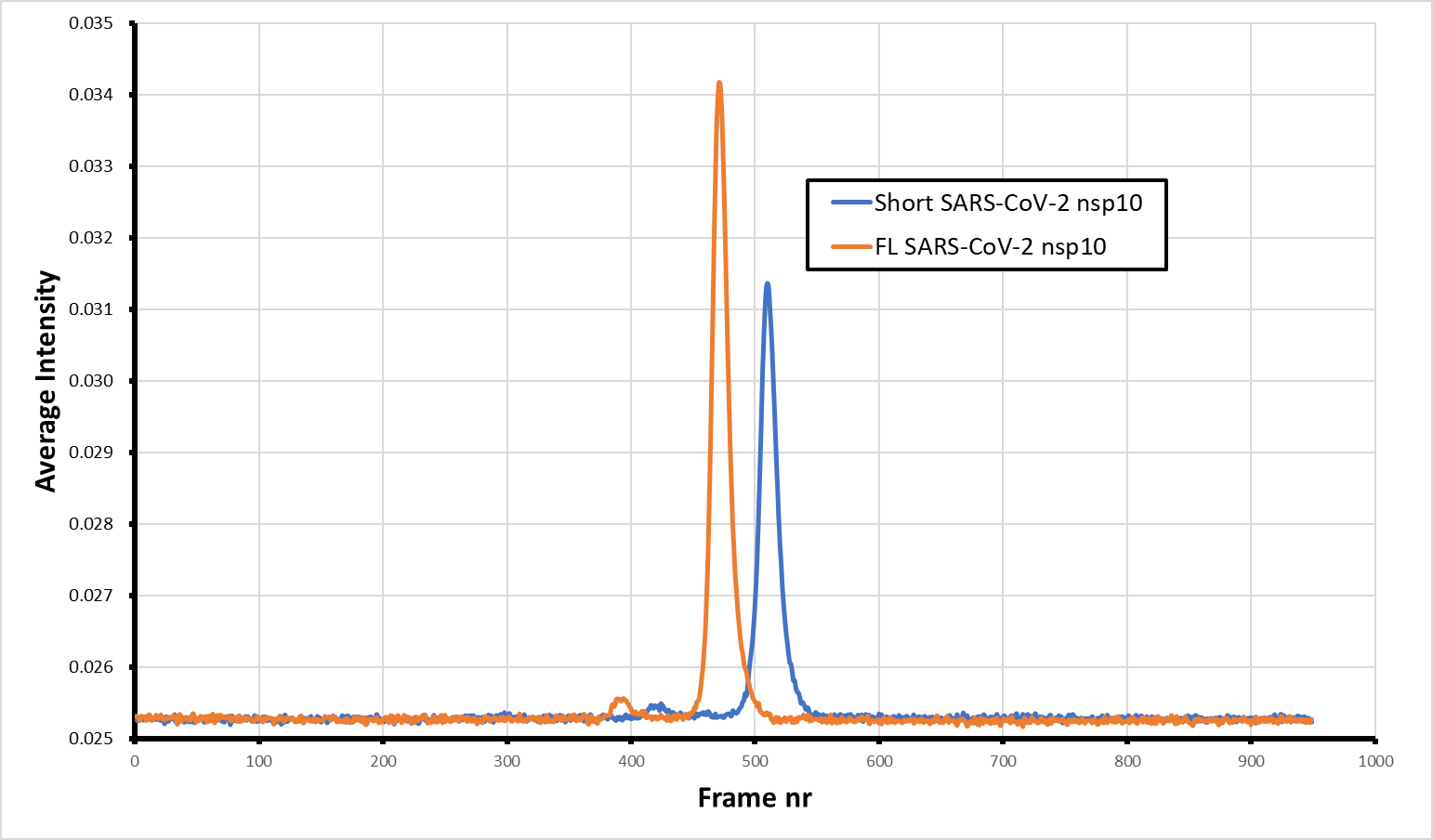

**Figure S4.** Kratky plots for the nsp10 proteins measured by SAXS as shown in Fig 2.

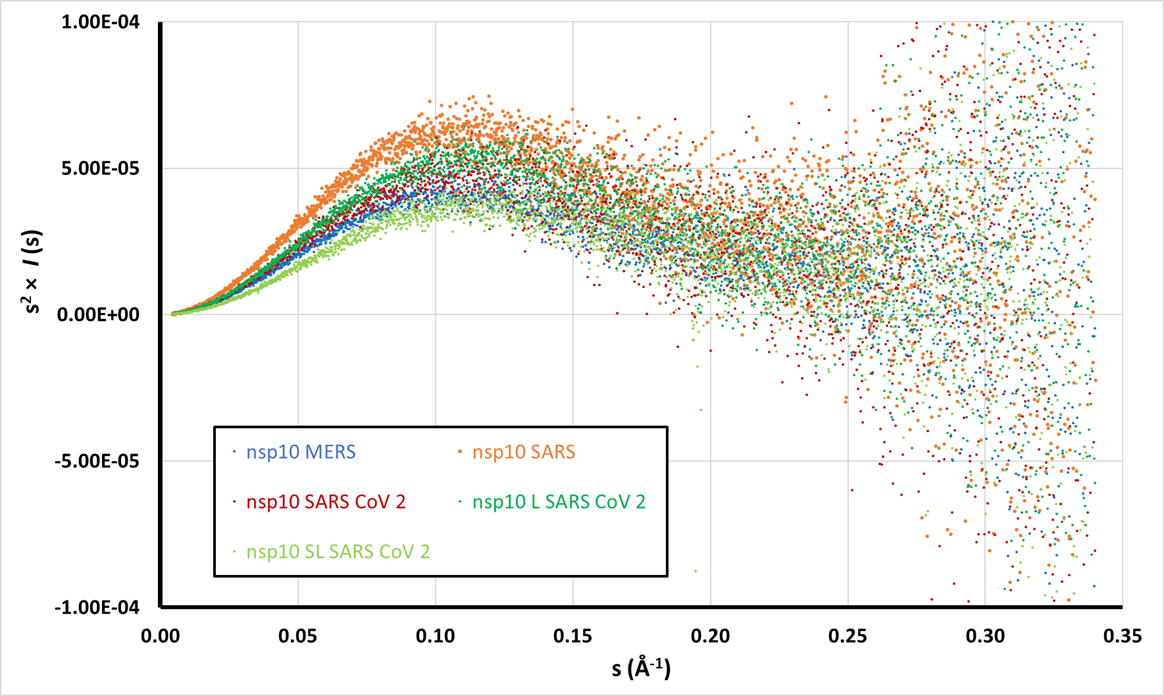
